# Evolutionary heritage shapes tree distributions along an Amazon-to-Andes elevation gradient

**DOI:** 10.1101/2020.06.11.143875

**Authors:** Andy R. Griffiths, Miles R. Silman, William Farfán Rios, Kenneth J. Feeley, Karina García Cabrera, Patrick Meir, Norma Salinas, Kyle G. Dexter

## Abstract

Understanding how evolutionary constraints shape the elevational distributions of tree lineages provides valuable insight into the future of tropical montane forests under global change. With narrow elevational ranges, high taxonomic turnover, frequent habitat specialisation, and exceptional levels of endemism, tropical montane forests and trees are predicted to be highly sensitive to environmental change. Using plot census data from a gradient traversing >3000 m in elevation on the Amazonian flank of the Peruvian Andes, we employ phylogenetic approaches to assess the influence of evolutionary heritage on distribution trends of trees at the genus level. We find that closely related lineages tend to occur at similar mean elevations, with sister genera pairs occurring a mean 254 m in elevation closer to each other than the mean elevational difference for all genera pairs. We also demonstrate phylogenetic clustering both above and below 1750 m a.s.l, corresponding roughly to the cloud-base ecotone. Belying these general trends, some lineages occur across many different elevations. However, these highly plastic lineages are not phylogenetically clustered. Overall, our findings suggest that tropical montane forests are home to unique tree lineage diversity, constrained by their evolutionary heritage and vulnerable to substantial losses under environmental changes, such as rising temperatures or an upward shift of the cloud base.

## 1. INTRODUCTION

Global environmental change urges investigation of potential evolutionary trends shaping biodiversity distribution and constraining the response of biota to novel environmental conditions (Lavergne et al., 2012; Christmas et al., 2016). Closely related lineages frequently occupy similar environments (Wiens & Graham, 2005, Holt, 2009), display similarities in functional characteristics (Felsenstein, 1985; Harvey & Pagel, 1991, Losos, 2008), and may respond comparably to environmental changes (Edwards and Donoghue, 2013). Understanding evolutionary influences on distribution patterns may provide a powerful aid in predicting the climate change response of unique and threatened systems such as tropical montane forests (TMFs). However, whether evolutionary heritage shapes distribution tendencies among TMF tree lineages is yet to be fully elucidated.

In keeping with montane environments globally, TMFs experience decreasing temperatures with increasing elevation (Humboldt and Bonpland, 1805; Schimper, 1903; Korner, 2007). Climatic trends, variation in biotic interactions (Hillyer et al., 2010), and topographic complexity engender globally exceptional levels of biodiversity within TMF (Myers et al., 2000; Hughes and Eastwood, 2006; Merckx et al., 2015). TMFs provide ecosystem services, regulating hydrological processes (Bruijnzeel et al., 2011) and influencing carbon and nutrient cycling (Girardin et al., 2010; Spracklen and Righelato, 2014; van de Weg et al., 2014). However, the response of TMFs to rapidly increasing temperatures (Pepin et al., 2015; Russell et al., 2017), precipitation regime changes (Hu and Riveros-Iregui, 2016) and other anthropogenic drivers remains poorly understood, with substantial potential for biodiversity losses (Feeley and Silman, 2010a; Feeley and Silman, 2010b).

Within TMFs, tropical montane cloud forests (TMCFs) are a fragile and enigmatic habitat, defined by persistent cloud immersion (Foster, 2001; Halladay, et al., 2012). Cloud immersion creates a complex, perhumid environment where epiphytes thrive (Bruinjzeel et al., 2011), but where light limitation can constrain the growth of other plants (Fahey et al., 2016). The lower elevation edge of TMCF, the cloud-base ecotone, marks a distinct transition within TMFs (Bruinjzeel, 2001; Fadrique et al., 2018) accompanied by changes in factors such as precipitation (Rapp & Silman, 2012), soil properties (Whitaker et al., 2014, Nottingham et al., 2015) and light availability (Fyllas et al., 2017.

The heterogeneous TMF environment combines with variation in climatic tolerances amongst biota to drive remarkable biodiversity (Richter et al., 2009) and distinct community level changes across elevations (Grubb and Whitmore, 1966; Hemp, 2006; Martin et al., 2011; Jankowski et al., 2013). Different climatic tolerances among species mean factors, such as temperature, filter the composition of communities across environments (Kraft et al., 2015). Such environmental filtering can have greater influence in more stressful environments (Chase, 2007), e.g. high elevation (Marx et al., 2017). If environmental filtering interacts with niche conservatism, the constraint of species’ environmental tolerances by their evolutionary history (Wiens and Donoghue, 2004; Wiens et al., 2010), then evolutionarily close lineages will be more likely to occur in similar environments (Cavender-Bares et al., 2009).

In montane landscapes, particularly at high elevations, evolutionary trends in distribution patterns are evident across a diversity of groups including microbes (Wang et al., 2012; Nottingham et al., 2018), ants (Machac et al., 2011), ferns (Kluge & Kessler, 2011), and alpine plants (Li et al., 2014). TMFs show significant dissimilarity in evolutionary composition of tree communities across different elevations (Ramirez et al., 2019). In the tropical Andes, distribution limits are manifest within certain tree genera. For example, *Weinmannia* (Cunoniaceae) and *Polylepis* (Rosaceae) tend to occur at higher elevations, while *Inga* (Fabaceae) and *Protium* (Burseraceae) tend to occur at lower elevations. However, the broader strength of evolutionary constraint on the elevational distribution of TMF tree lineages remains unclear.

Rapid environment change forces biodiversity to adapt, acclimate, migrate, or face extinction (Aitken et al., 2008; Feeley et al., 2012). With rates of upslope migration insufficient for most taxa to track predicted temperature changes (Feeley et al., 2011; Fadrique et al., 2018), some TMF lineages may endure through acclimation or adaptation. High taxonomic turnover (Bach et al., 2007; Baldeck et al., 2016) and narrow elevational range sizes (Terborgh, 1977; Lieberman et al., 1996; Perez et al., 2016) are predominant within TMF. However, a few lineages, like the genera *Miconia* (Melastomataceae) and *Meliosma* (Sabiaceae), occupy wide elevational ranges, encompassing broad climatic variation. Such labile taxa may possess an adaptive potential that is advantageous in responding to climate change.

Evolutionary accessibility to potential adaptations may be phylogenetically constrained (Edwards and Donoghue, 2013). For example, C4 photosynthesis in grasses has only evolved in the PACMAD clade, a lineage possessing certain enabling traits (Christin et al., 2013). Similarly, constraints on adaptation to freezing conditions, combined with niche conservatism, may explain why the expansion of angiosperm lineages to temperate regions is phylogenetically biased (Wiens and Donoghue, 2004; Mittelbach et al., 2007; Donoghue, 2008; Zanne et al., 2014). If tolerance of variation in for example, temperature, moisture regime, or cloudiness is phylogenetically clustered within only a few TMF tree lineages, the remaining TMF diversity may be threatened by environmental change.

Testing for phylogenetic signal (PS) provides a simple means of quantifying correlations between evolutionary history and characteristics such as elevational distribution. A statistical measure of the non-independence of trait values of taxa due to evolutionary relatedness, PS quantifies the tendency for closely related taxa to resemble each other more than they resemble taxa drawn randomly from a phylogeny (Felsenstein, 1985; Losos, 2008; Revell, Harmon, and Collar, 2008). Evidence of PS has previously been shown for diverse characteristics in tropical trees, from mean range size and abundance (Dexter and Chave, 2016) to wood density, tree size, and mortality rates (Coelho de Souza et al., 2016).

Based on an Amazon-to-Andes elevation gradient, this study investigates potential evolutionary constraints on elevational distribution and response to environmental change within TMF tree lineages. A temporally-calibrated, genus-level phylogeny is generated – covering the breadth of vascular plant diversity, from angiosperms and gymnosperms to tree ferns. Using this phylogeny, we test for evolutionary patterns underlying elevational distribution trends and the influence of the cloud-base ecotone. Specifically, we consider two core hypotheses: 1) closely related genera occupy similar elevations, and 2) genera displaying environmental lability are phylogenetically clustered.

## 2. METHODS

### 1. Study site

A plot network spanning a 425 to 3625 m a.s.l Amazon-Andes elevation gradient centred on Kosñipata valley, both in and near Manu National Park, south-eastern Peru (Figure 1). Established by the Andes Biodiversity and Ecosystem Research Group (ABERG: www.andesconservation.org), plots are subject to regular re-censusing and ongoing multidisciplinary research (Malhi et al., 2010; Malhi, 2017). The gradient encompasses broad habitat and environmental variation, from lowland/sub-montane forests below 800 m a.s.l, up to the montane forest-puna grassland transition ∼3400 m a.s.l (Girardin et al., 2010). Mean annual temperature decreases from ∼24°C at low elevations, to ∼9°C at higher elevation (Malhi et al., 2017). Mean total annual precipitation displays a hump shaped trend across the gradient, ranging from ∼3000 mm yr^1^ at low elevations, ∼5000 mm yr^1^ at mid elevations, and ∼1000 mm yr^1^ at high elevations (Rapp and Silman, 2012). Frequent cloud immersion, characteristic of TMCF, occurs above 1500-2000 m a.s.l (Girardin et al., 2010; Rapp and Silman, 2012), reaching peak frequency between 2000-3500 m a.s.l (Halladay et al., 2012). Paleozoic meta-sedimentary mudstone dominates geologically, with granite intrusions between 1500-2020 m a.s.l (Nottingham et al., 2018). Significant soil character changes occur ∼1000 and ∼2000-2500 m a.s.l (Nottingham et al., 208)

**Figure 1.**
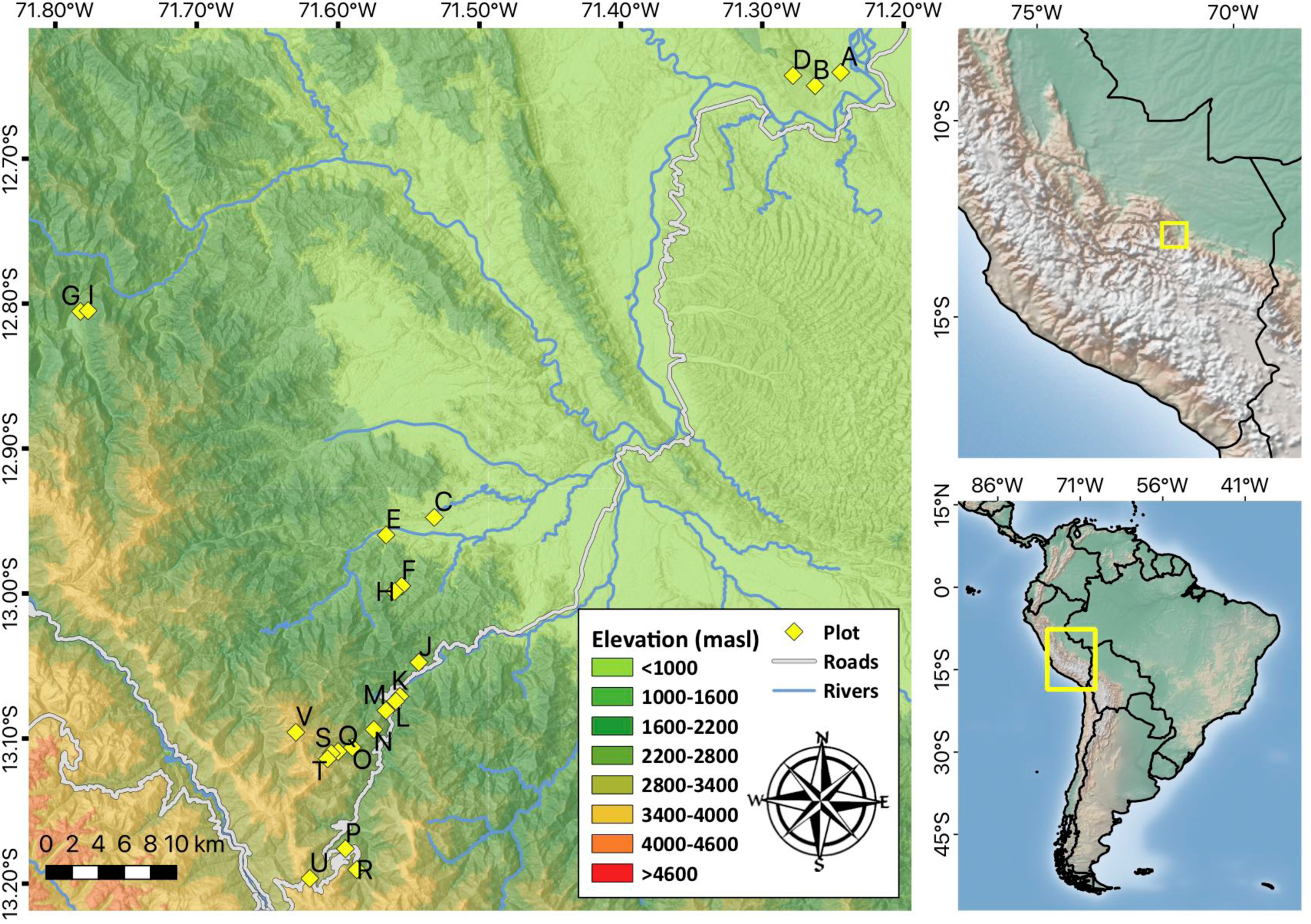
Location of 23 plots along an elevation gradient on the Amazonian flank of the south-eastern Peruvian Andes. Yellow diamonds indicate location of plots. Letters relate to the following plots and elevations (m a.s.l): A: PAN-01 (425), B: PAN-02 (595), C: TON-01 (800), D: PAN-03 (850), E: TON-02 (1000), F: SAI-02 (1250), G: CAL-02 (1250), H: SAI-02 (1500), I: CAL-01 (1500), J: SPD-02 (1500), K: SPD-01 (1750), L: TRU-08 (1800), M: TRU-07 (2000), N: TRU-06 (2250), O: TRU-05 (2500), P: TRU-04 (2750), Q: ESP-01 (2890), R: TRU-03 (3000), S: WAY-01 (3000), T: TRU-02 (3250), U: TRU-01 (3450), V: ACJ-01 (3537), W: APK-01 (3625).

### 2. Plot inventory & phylogeny

This study utilises the latest inventory data for woody stems >10 cm diameter at breast height (1.3m above the ground; DBH) growing in 23 1-hectare plots within the ABERG plot network (ABERG: www.andesconservation.org). 301 plant genera were inventoried across all plots. Sequences for rbcL and matK plastid genes were obtained from the GenBank database (www.ncbi.nlm.nih.gov/genbank/; Benson et al., 2017), for 267 and 259 of these genera respectively (251 genera with both rbcL and matK sequences). Where possible, sequences were used for geographically proximate representatives of each genus. Sequences were aligned on the MAFFT v7 online service (https://mafft.cbrc.jp; Katoh et al., 2017). Manual alignment checking and trimming of sequence ends where most genera lack data, was carried out in Mesquite v3.6 (Maddison and Maddison, 2018). Since rbcL and matK are chloroplast markers and therefore do not experience recombination, sequences were concatenated prior to phylogeny estimation.

A maximum likelihood phylogeny was estimated for the 275 genera in RAxML-HPC2 v8.2.10 (Stamatakis, 2014), with rapid bootstrapping (100 iterations), executed on the CIPRES web server (www.phylo.org; Miller, Pfeiffer and Schwartz, 2010) under default settings including a General Time Reversible (GTR) + Gamma (G) model of sequence evolution. Family level relationships were constrained using the ‘R20160415.new’ megatree (Gastauer and Meira-Neto, 2017), based on the APG IV topology (Chase et al., 2016). Phylogeny temporal calibration was conducted utilising penalised likelihood methods in treePL (Smith and O’Meara, 2012) with secondary calibrations on 59 of 275 internal nodes, based on age estimates in Magallon (2015) for angiosperms; Silvestro et al (2015) for further angiosperms and gymnosperms; Lu et al (2014) for Podocarpaceae; and Korall and Pryer (2014) for Cyatheaceae.

Patterns often vary across phylogenetic scales (Graham et al., 2018). As such, both our full phylogeny (275 genera) and a subset comprising only angiosperm genera (269 genera) were analysed to investigate the consistency of potential phylogenetic trends. Our full phylogeny encompasses a phylogenetic depth much older than the angiosperms with gymnosperms and tree fern lineages occurring on long branches, which when included in analyses may mask phylogenetic trends at the angiosperm scale (Kembel and Hubbell 2006, Honorio-Coronado et al. 2015).

### 3. Elevational distribution trends

To test for evolutionary patterns, elevational characteristics of genera were calculated and mapped onto our phylogeny. To quantify similarity of elevational distribution among close relatives, we used abundance weighted mean elevations of genera, based on numbers of individuals per genus within plots. We used several metrics to assess the capacity for lineages to respond evolutionarily to novel environmental conditions, including measures of lability for elevational preference: 1) a metric of temporal elevational shifts in the distribution of genera, based on their annualised change in basal-area weighted mean elevation, quantified on this transect by Feeley et al (2011) over a four year period for 35 genera within our phylogeny. 2) Elevational range breadth for all 275 genera in the phylogeny, measured as the 95% quantiles of occurrence for each genus on the gradient. 3) Coefficient of variation (CV) for mean elevation of species within a genus, for the 148 genera with more than one species on the transect (the other 127 genera are monotypic across sampling sites). Both elevational range breadth and CV for mean elevation are indicative of broad environmental tolerances. High values in both measures suggest a genus occupies a breadth of environmental variation and may therefore be better able to tolerate climatic changes. A metric of change over time may better represent the potential of genera to respond to climatic changes, but is only quantified for 35 genera in our phylogeny (Feeley et al., 2011). We therefore assessed if our other measures of evolutionary lability (elevational range breadth and CV of mean elevation), available for many more genera, correlate with annual change and may therefore act as proxies.

We estimated phylogenetic signal for genera elevational characteristics using Pagel’s λ (Pagel, 1999; Freckleton et al., 2002). Based on a comparison of tree branch length transformations, λ contrasts variance in observed trait values against expected trait variance under a Brownian motion (BM) model of evolution. Under a BM model, trait values evolve following a stochastic random walk trajectory, with expected trait divergence across each node in the phylogeny being proportional to the phylogenetic depth, or age, of the node. This random walk results in a linear increase in variance with time, and therefore, variance and covariance of trait values between lineages proportional to phylogenetic branch length. Values of λ around 0 indicate no phylogenetic signal. Values of λ around 1 indicate strong phylogenetic signal, matching that expected under a BM model of evolution. Values of λ between 0 and 1 indicate intermediate levels of phylogenetic signal. In order to test whether results display metric dependency, we also calculated phylogenetic signal using Blomberg’s K (Blomberg et al., 2003).

Following logic similar to the mid-domain effect on species richness (Colwell et al., 2000), if phylogenetic signal exists for elevational range breadth, any phylogenetic signal for mean elevation could be an artefact driven by constraint of mean elevations for widely ranged genera to intermediate elevations. To test for this potential artefact we ran null simulations, shifting the ranges of genera randomly up and down while ensuring ranges remained with the gradient limits. We biased genera placements to retain the observed distribution pattern or range mid-points, including a higher frequency of mid-points at lower elevations. We compared the mean phylogenetic signal for genera range mid-point from 1000 null simulations with our observed phylogenetic signal for abundance weighted mean elevation. Range mid-point is strongly correlated with abundance weighted mean elevations (R^2^ = 0.93) and give similar measures of phylogenetic signal in the observed dataset.

### 4. Cloud-base ecotone

We additionally investigated the influence of the cloud-base ecotone on elevational distribution patterns. Along our gradient, the cloud-base occurs consistently between approximately 1500-2000 m a.s.l (Girardin et al., 2010; Rapp and Silman, 2012). We used the mid-point of this range (1750 m a.s.l) as the cloud-base elevation in our analysis. The robustness of this cloud-base approximation was tested with a hierarchical cluster analysis using Bray-Curtis dissimilarity indices of plots based on their species composition. We then categorised genera based on their distribution relative to the cloud base. Three distribution categories were assigned 1) only above cloud-base (≥1750 m a.s.l; n = 33), 2) only below cloud-base (≤1749 m a.s.l; n = 161), or 3) occurring across the cloud-base (both <1749 m a.s.l and >1750 m a.s.l; n = 81). We estimated phylogenetic signal for each category using the D statistic for discrete characters (Fritz and Purvis, 2010). D is based on the sum of sister clade differences. Running opposite to Pagel’s λ values, a D value of 1 indicates no phylogenetic signal, and a D value of 0 indicates phylogenetic signal equivalent to that expected under a BM model of evolution. Values < 0 and > 1 are possible. The observed value is then assessed for significance against the expected value, generated from simulations (n = 5000) based on an absence of phylogenetic dependency, and phylogenetic structure based on a BM model of evolution.

## 3. RESULTS

### 1. Elevational distribution trends

Abundance weighted mean elevation shows high and significant phylogenetic signal at the genus level (λ = 0.81, p < 0.001), though slightly less than expected under a Brownian motion model of evolution. Phylogenetic signal is also observed when considering only angiosperm lineages (λ = 0.62, p < 0.001), suggesting the effect is consistent across the phylogenetic scales of our analysis. The mean (λ = 0.02, sd = 0.07) and maximum (λ = 0.52) phylogenetic signal generated by null simulations of genera elevational range mid-points suggest our observed λ value is not simply an artefact akin to the mid-domain effect (Colwell et al. 2000).

Significant phylogenetic signal for mean elevation is driven by high and low mean elevation values across a number of lineages (Figure 2a). High mean elevation values occur frequently across the Asterids, with the notable exceptions of the Apocynaceae, Rubiaceae, Sapotaceae, and Lecythidaceae which tend towards lower mean elevations. In contrast, low mean elevation values are more dominant within the Rosids; strongly so among the Malpighiales, Fabaceae, and Malvaceae. Exceptional among Rosid lineages, the Oxalidales and Melastomataceae tend towards high mean elevations. Arecaceae, the sole Monocot lineage in the phylogeny displays a low mean elevation pattern. The Magnoliids are largely split between a low mean elevation trend within the Annonaceae and Myristicaceae, and a mid-elevation mean within the Lauraceae. Beyond the angiosperms, the Podocarpaceae and Cyatheales lineages also display largely mid-elevation means. The difference between the mean elevation of genera is 252 m lower for sister genera in the phylogeny (n = 83, mean = 504 m, sd = 506 m) than it is between all pairs of genera (n = 37675, mean = 756 m, sd = 647 m).

**Figure 2.**
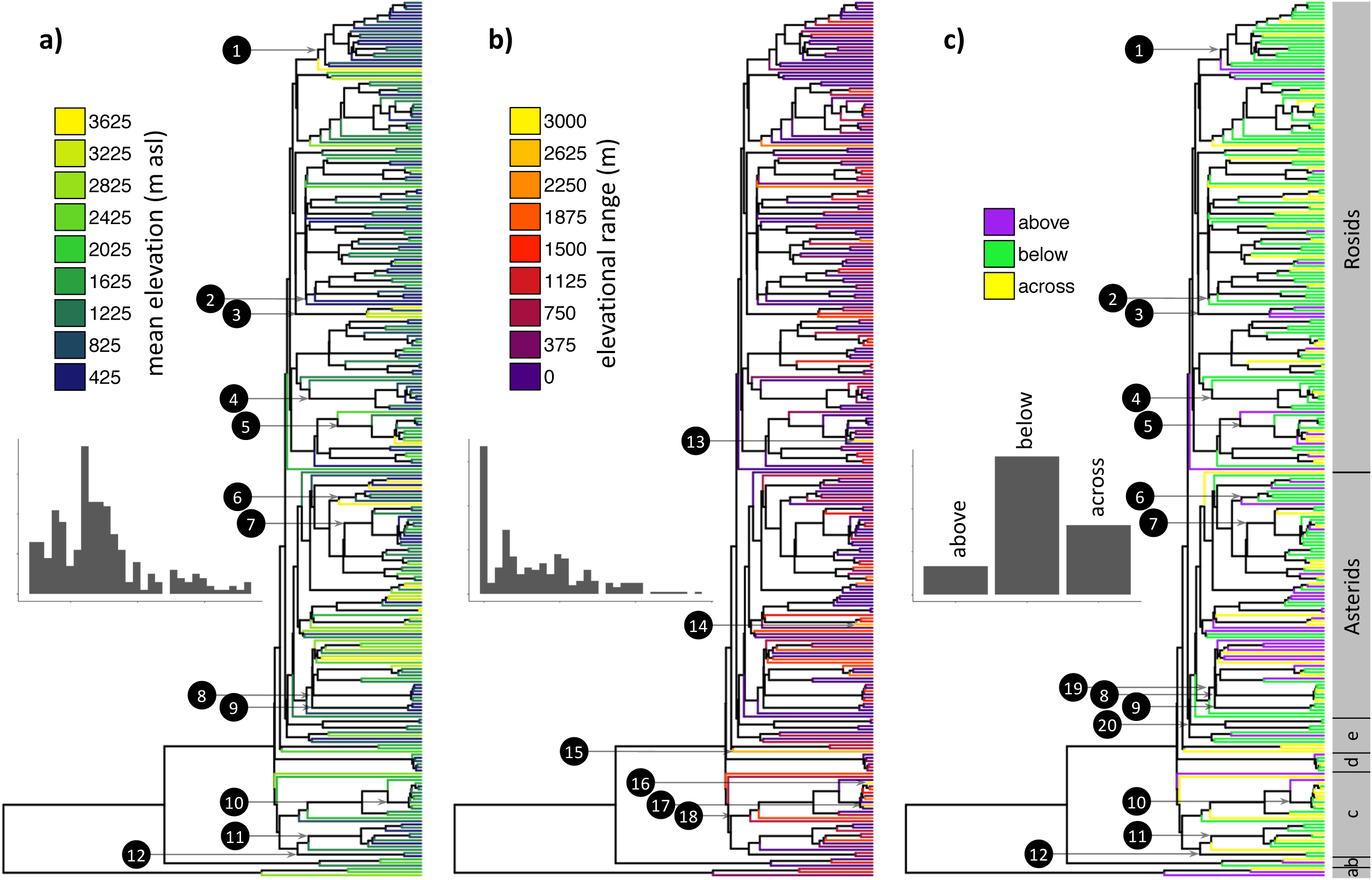
Phylogeny of 275 tree genera with terminal branches coloured according to: a) abundance weighted mean elevation; b) elevational range size; and c) distribution relative to the cloud-base ecotone. There is significant phylogenetic signal for a) mean elevation (λ = 0.81, p < 0.001) and c) distribution solely above (D = 0.31, p < 0.001) or solely below (D = 0.69, p < 0.001) the cloud-base ecotone. There is no significant phylogenetic signal for b) genera elevational range size (λ = 0.36, p = 0.09) or c) distribution across the cloud base-ecotone (D = 0.9, p = 0.16). Major clades are indicated in grey bar to the right side: a = tree ferns (Cyatheales), b = gymnosperms, c = Magnoliids and *Hedyosmum*, d = Monocots, e = basal Eudicots. Numbered nodes indicate branch stems of lineages mentioned in the main text: 1 = Fabaceae, 2 = Malpighiales, 3 = Oxalidales, 4 = Malvales, 5 = Melastomataceae, 6 = Apocynaceae, 7 = Rubiaceae, 8 = Sapotaceae, 9 = Lecythiadaceae, 10 = Lauraceae, 11 = Annonaceae, 12 = Myristicaceae, 13 = Miconia, 14 = Schefflera, 15 = Meliosma, 16 = Ocotea, 17 = Persea, 18 = Laurales, 19 = Ericales, 20 = Caryophyllales. Inset bar plots display frequency distribution of values for each variable.

The annual change in the mean elevation of genera, weighted by relative basal area, is positively correlated with both elevational range breadth of genera (τ = 0.46, p < 0.001; Figure 3a) and the coefficient of variation for mean elevation of species within genera (τ = 0.28, p = 0.01; Figure 3b). These correlations suggest elevational range breadth and coefficient of variation for mean elevation may be acceptable proxy measures for the adaptive potential of genera to change their elevational distribution.

**Figure 3.**
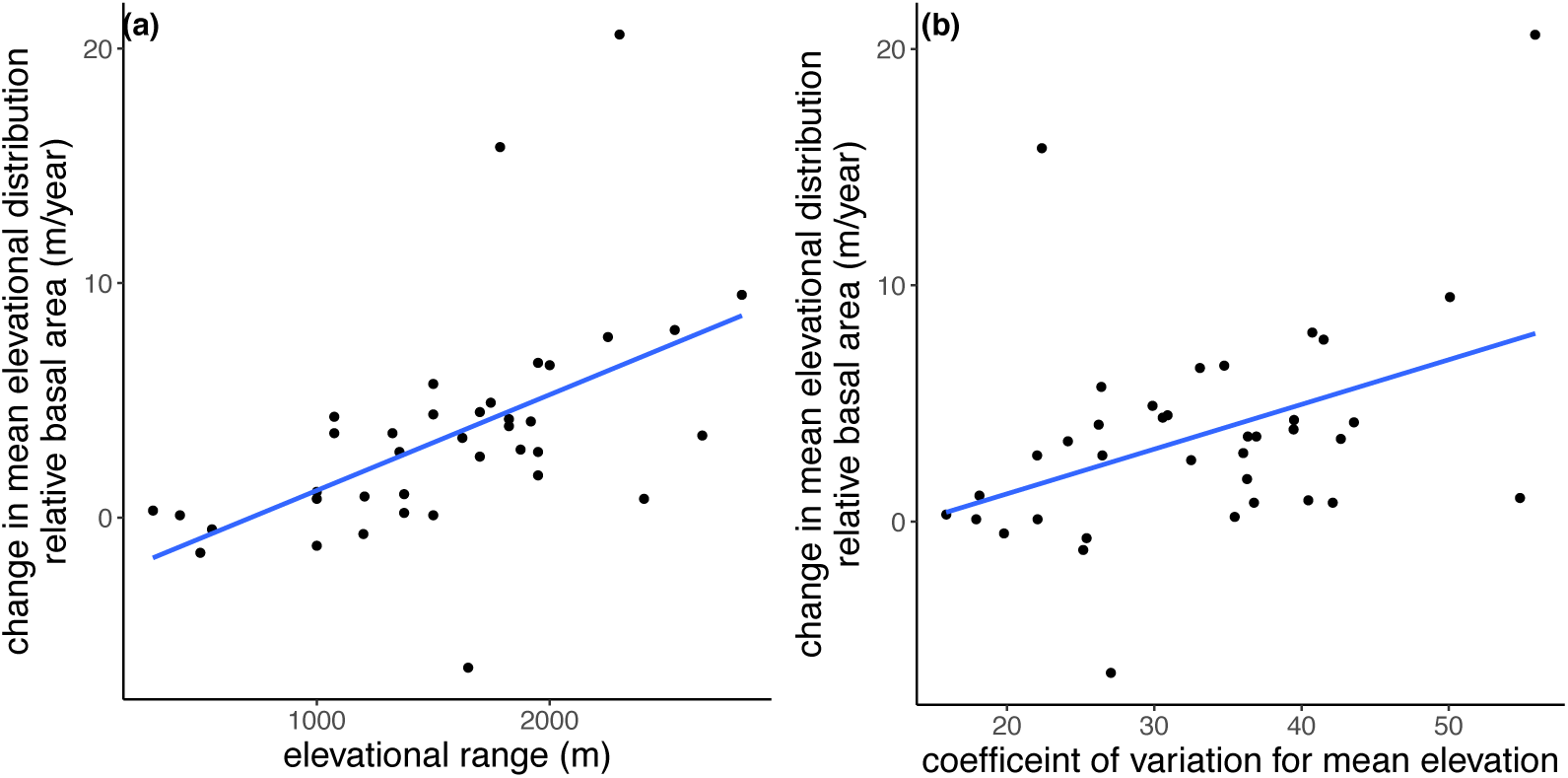
The annual change in mean elevation of genera, weighted by relative basal area, is positively correlated with a) the elevational range breadth of genera (τ = 0.46, p < 0.001), and b) the coefficient of variation for mean elevation of species within genera (τ = 0.28, p = 0.01). Points represent genera. Correlations based on Kendall’s tau coefficient. Blue lines are derived from linear regression.

There is no phylogenetic signal for annual change in the mean elevation of genera, weighted by relative basal area (λ < 0.001, p = 1, n = 35). Similarly, no significant phylogenetic signal is evident for elevational range size of genera (λ = 0.36, p = 0.09, n = 275; Figure 2b), or the coefficient of variation of species mean elevations within genera (λ = 0.00007, p = 1, n = 148). Analysing angiosperm lineages alone reveals significant values for elevational range size (λ = 0.2, p = 0.01, n = 269), though this is not consistent across metrics (Blomberg’s K = 0.2, p = 0.14).

### 2. Cloud-base ecotone

A hierarchical cluster analysis identifies clear dissimilarity in species composition across plots. Though edaphic and topographic variations between plots must be noted, mid-elevations appear to be a clear point of species turnover, with all plots at 1800 m a.s.l and above more similar to each other in species composition than they are to all plots at 1750 m a.s.l and below, and vice-versa (Figure 4). This pattern is driven by the fact that more species reach the limit of their elevational distribution before 1750 m a.s.l than at other elevations. These patterns lend support to our approximation of 1750 m as the elevation for the cloud-base ecotone and its relevance as an area of ecological transition.

**Figure 4.**
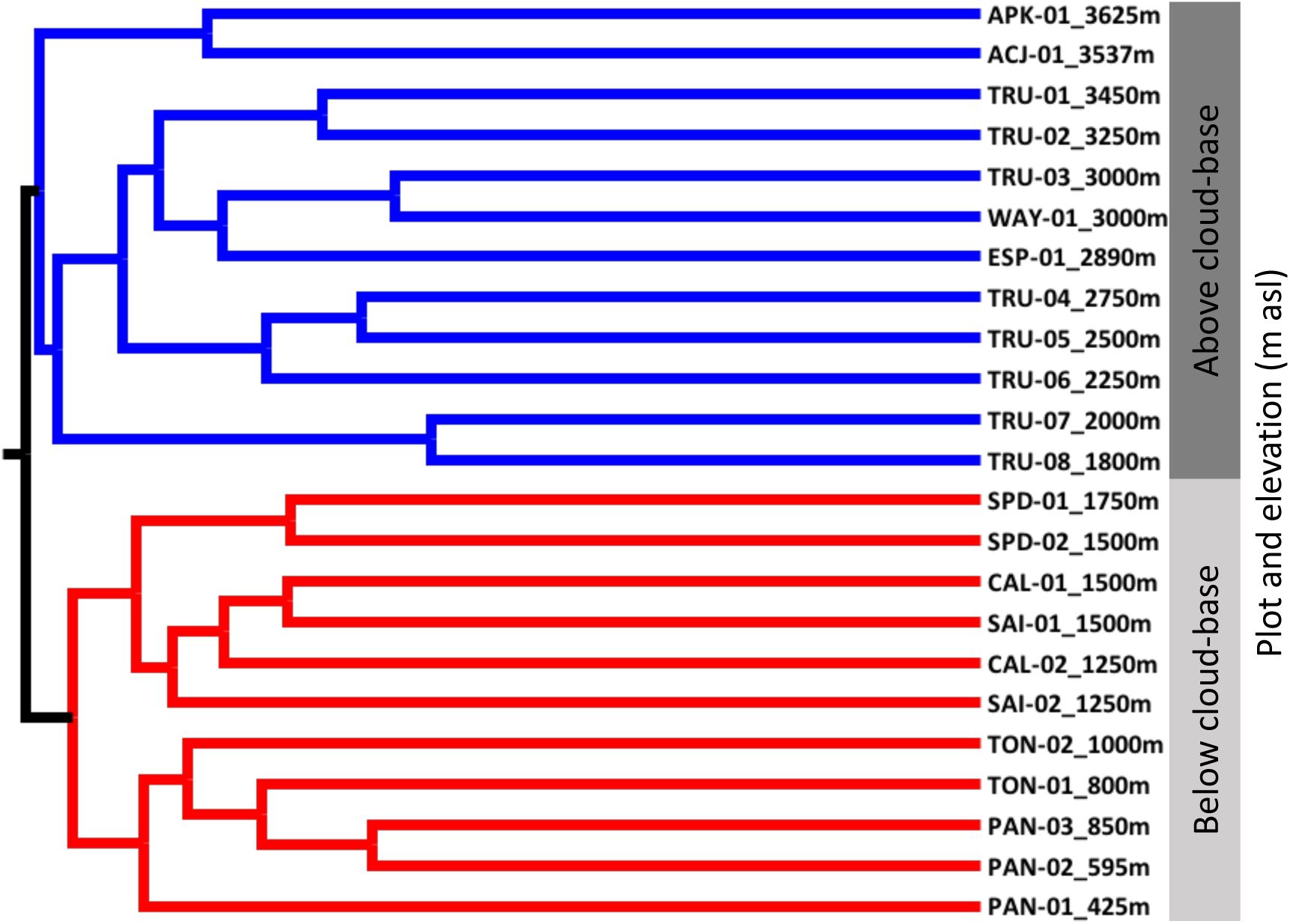
Dendrogram generated by a hierarchical cluster analysis based on Bray-Curtis dissimilarity indices, illustrating the main areas of taxonomic turnover across the elevation gradient. Species compositions in all plots at or above 1800 m a.s.l (indicated by blue branches) are more similar to each other than species compositions in all plots at or below 1750m (indicated by red branches) and vice-versa.

Genera distributed solely above the cloud-base ecotone (n = 33), are more significantly clustered in the phylogeny than would be expected under a model of random phylogenetic structure (D = 0.31, p < 0.001; Figure 2c). Further, the observed phylogenetic signal is not significantly different than expected under a Brownian model of evolution (p = 0.12). An above cloud-base distribution is more frequent among Asterid genera, notably so for a clade within the Ericales. Genera occurring at elevations solely below the cloud-base ecotone (n = 161) are also more significantly clustered in the phylogeny than expected under a random phylogenetic structure (D = 0.69, p = 0.001; Figure 2c), though less than expected under a Brownian model of evolution (p < 0.001). Below cloud-base distribution is common among Rosid lineages, notably genera within the Fabaceae, Malvaceae, and Malpighiales. While there is also a strong trend within the Apocynanaceae, Sapotaceae, Lecythidaceae, Annonaceae, and Caryophyllales for genera with below cloud-base distributions. Those genera occurring across the cloud-base ecotone, i.e. those that show lability in occurrence across this environmental threshold (n = 81), are not significantly clustered in the phylogeny (D = 0.9, p = 0.16; Figure 2c). These results are also consistent when only angiosperm lineages are considered in analyses.

## 4. DISCUSSION

We find clear phylogenetic signal for the mean elevational occurrence of genera, suggesting that evolutionary heritage strongly influences elevational distributions of tree genera within tropical montane forests. The observed phylogenetic signal is higher than that measured previously for a number of tree functional characteristics (Coelho de Souza et al., 2016; Baraloto et al., 2012). Closely related tree genera tend to occupy similar mean elevations, clustering either above or below the cloud-base ecotone and its associated environmental changes. Further, those genera that occur above the cloud base show stronger phylogenetic clustering than those below the cloud base. While the general pattern is for narrow elevational ranges of genera as a whole, some genera appear able to escape this evolutionary constraint, occupying large elevational ranges and crossing the cloud-base ecotone. These more broadly distributed genera are not phylogenetically clustered, but rather seem to arise randomly across the breadth of vascular plant lineages represented in our phylogeny.

That closely related genera tend to occupy similar mean elevations is evidence that evolutionary heritage influences biodiversity distribution across the heterogeneous environment of tropical montane forests. This observed trend, in combination with high taxonomic turnover (Mahli et al., 2010; Jankowski et al., 2013; Baldeck et al., 2016) and narrow elevational ranges (Perez et al., 2016), is consistent with the prediction of niche conservatism that it tends to be hard for lineages to evolve environmental tolerances that differ markedly from those of their evolutionary ancestors (Donoghue, 2008; Wiens et al., 2010). Our genus level observations build upon Gentry’s (1988) convincing demonstration of elevational change in dominant lineages across tropical montane forests at the family level. Gentry (1988) noted for example, the replacement of lowland dominant Fabaceae by Lauraceae at intermediate elevations, and the dominance of Asterid families such as Asteraceae and Rubiacae at higher elevation. These trends are evident in our observations, while the overrepresentation of Asterids lineages at high elevation has also been observed elsewhere (Hemp, 2006; Molina-Venegas et al., 2020) and deserves further exploration.

The shifting evolutionary composition of communities across elevation is further reinforced by the observed phylogenetic clustering of closely related genera solely above, and solely below the cloud-base ecotone. Associated with sharp climatic changes, such as reduced solar radiation and increased occult precipitation, the cloud-base ecotone may represent an important environmental barrier, constraining the movement of lineages between contrasting environments (Pounds et al., 1999; Fadrique et al., 2018). A cluster analysis revealing strong dissimilarity in species composition between plots above versus below the cloud-base, also shown by Jankowski et al. (2013), further suggests that this ecotone is an area of significant floristic transition (Figure 4). Higher phylogenetic signal for lineages distributed solely above the cloud-base ecotone compared to those solely below, suggests that clustering of lineages, niche conservatism, and environmental filtering, may be stronger drivers of distribution patterns within TMCF. Frequent cloud immersion presents ecological challenges which may engender particular adaptations, such as foliar water uptake (Eller et al., 2013; Goldsmith et al., 2013), and promote the evolution of a unique floral diversity above the cloud-base ecotone.

Any cloud-base effect is likely to act in concert with other factors. For example, ∼1700 m a.s.l may represent the upper temperature limit for many tropical forest tree species which have tracked to higher elevations after evolving under the cooler lowland conditions of the Pleistocene (Silman, 2007). Additionally, the geologically recent major uplift of the Andes, and relative lack of high elevations until <10 mya (Garzione et al., 2008; Leier et al., 2013; Sundell et al., 2019), likely had major impact on the biogeographic history of the South American flora (Antonelli et al., 2009). However, evidence is increasingly suggesting a genuine cloud-base effect. For example, phylogenetic discontinuity between cloud forest and lower montane forest occur in Africa on the slopes of Mt Kilimanjaro (Molina-Venegas et al., 2020). Though further comparisons between TMFs are necessary, it seems likely that TMCFs contain a unique and important component of biodiversity. Along our gradient, lineages with a clear temperate affinity, such as *Alnus* (Betulaceae) and *Prunus* (Rosaceae), are observed more frequently at middle and upper elevations yet these lineages do not dominate. Rather the tree assemblages of tropical montane cloud forests appear to contain substantial unique evolutionary diversity.

It is important to note that while the characteristics analysed show significant phylogenetic signal, this signal is generally less than that expected under a Brownian motion (BM) model of evolution, where λ values would be close to 1 and D values close to 0. This may be the result of divergent selection amongst closely related taxa and/or convergent evolution across distant relatives. On the other hand, a simple BM model may not accurately describe genus-level distribution changes over time. For example, a simple BM model does not account for variation in rates of evolution over time or among lineages. Different models of evolution are possible, but our goal was simply to identify the existence of phylogenetic signal and not to test any specific underlying evolutionary mechanism, given the genus-level nature of our phylogeny. As such, a BM model can provide insight into the evolutionary trends influencing elevational patterns such as high taxonomic turnover and narrow elevational range occupancy.

Most genera evidently occupy relatively narrow elevational distributions. However, a few genera, such as *Miconia* (Melastomataceae), *Meliosma* (Sabiaceae), *Ocotea* (Lauraceae), *Persea* (Lauraceae) and *Schefflera* (Araliaceae), seem able to escape the constraints of evolutionary heritage and occupy large elevational ranges, as well as cross the ecotonal transition of the cloud-base (Figure 2b-c). In addition to occupying broad elevational ranges, genera such as *Miconia, Persea*, and *Schefflera*, are among those that show significantly greater upslope shifts in mean elevation than tree genera as a whole (Feeley et al., 2011). In the cases of *Miconia* and *Schefflera*, rates of elevational change have actually kept pace with predicted temperature increases, which contrasts with most other tree genera that are lagging in their responses to temperature increases (Feeley et al., 2011, Malhi et al., 2009, Urrutia and Vuille, 2009). Such trends in specific genera, along with the more general correlation observed between elevational range size and rate of elevational distribution change (Figure 3a-b), reinforce the suggestion that occupancy of a broad elevational range may be associated with a greater lability of response to the pressures of a changing climate. In any case, our findings reveal no phylogenetic signal for elevational range size (Figure 2b) or annual rate of elevational distribution change, demonstrating that characteristics such as broad elevational ranges, or trends of upslope distribution change, are not phylogenetically clustered among closely related genera. Rather, such genera come from lineages distributed across the breadth of the vascular plant phylogeny.

The observed random phylogenetic pattern for elevational range size provides an interesting contrast to research revealing clear phylogenetic signal for geographic range size across Amazonian tree lineages (Dexter and Chave, 2016). However, the environmental drivers of elevational range sizes, more closely linked to abiotic tolerances (Janzen, 1967; Ghalambor et al., 2006), are likely to be different to those operating on range size in the tropical lowlands where biotic effects are thought to be stronger. While phylogenetic signal is not evident for elevational range size across the breadth of vascular plant genera considered in this analysis, there appears to be a trend for broad elevation ranges in a few lineages, notably the Laurales (Figure 2b). Such lineages may drive the marginally significant phylogenetic signal observed for elevational range size when only angiosperms are included in the analysis.

Although lability of response to environmental change, indicated by occupancy of a broad elevational range, is not clearly constrained within certain evolutionary lineages, the majority of lineages nonetheless occupy narrow elevational ranges, and the timescale necessary for evolutionary adaptation within most tree lineages may be incompatible with the current rapid rate of environmental change (Feeley et al., 2011; Pepin et al., 2015; Russell et al., 2017). Clustering of closely related genera around similar mean elevations may suggest that climatic trends, such as rising temperatures, will have unequal impacts across lineages. Lowland lineages, already occupying broad distributions across the Amazon, may find that amenable environmental conditions become available on higher ground. Meanwhile those few lineages already occupying broad elevational distributions may find themselves at a competitive advantage in terms of tolerating changing conditions. However, TMF lineages, and the evolutionary diversity constrained to mid and high elevations may be at risk. As climate conditions track up mountain slopes, the area of land amenable to TMF lineages will be reduced in size or disappear completely. At the same time TMF lineages may be squeezed from below by increasing competition as lowland lineages migrate upslope (Colwell et al., 2008; Feeley et al., 2011). Among TMF lineages, those clustered solely above the cloud-base ecotone may be most vulnerable. Many TMCF tree lineages display adaptation to the conditions associated with frequent cloud immersion, such as high foliar water uptake, making them specialised and at risk under a changing climate (Eller et al., 2016). With lineages constrained by evolutionary heritage to narrow elevational distributions and particular habitats, climatic changes such as a decline in frequency of cloud immersion and lifting of the cloud base (Still et al., 1999; Helmer et al., 2019), may fundamentally alter the tropical montane environment and result in large population reductions and potential extinctions among the TMF biota. Phylogenetic clustering in the elevational distribution of TMF tree lineages means any extinctions may lead to a disproportionate loss of evolutionary history, a risk which is particularly stark for specialised lineages constrained to TMCF.

A degree of perspective must be given to interpretation of phylogenetic analyses at the genus-level. Genera may encompass substantial variation among their constituent species and species-level patterns may influence genus-level trends. For example, along our gradient the range size of genera is correlated with the mean range size of their constituent species, although the latter only explains 16% of the variation in the former. In addition, species-level analyses may reveal divergent patterns at a finer scale, though DNA sequence data are not yet available for an analysis representing the breadth of lineages we can consider at the genus-level. Further, focusing on higher taxonomic levels, such as genera, can be a valuable means to understanding deeper evolutionary trends. Additionally, most species in this data set are only recorded in a single plot and therefore, given elevational intervals up to 250 m between plots, quantification of their elevational distribution would have limited precision. A genus level analysis is also advantageous in that it minimises potential errors created by any individuals not reliably identified to species. Nevertheless, future analyses of lineage specific, species-level phylogenetic trends across elevation, particularly focusing on functional characteristics, would further develop our understanding of the mechanisms driving elevational distribution patterns.

Overall, our study illustrates that by utilising phylogenetic approaches we can better understand how evolutionary heritage, and the tendency of close relatives to share similar ecological and functional characteristics, influences lineage distribution patterns across different environments. In particular, our analyses draw out the ecological significance of environmental transition zones, such as the cloud-base ecotone, showing that such transitions can coincide with significant phylogenetic community turnover. Further, by demonstrating clustering of evolutionary lineages at similar elevations, we provide valuable insight into the potential impact rapid environmental changes may have on the unique and vulnerable evolutionary diversity of tropical montane forests in general, and tropical montane cloud forests in particular.

## Author Contributions

ARG and KGD conceived the analyses. MS, WFR and PM designed and established the plot network. KJF, WFR, KCG, and NS contributed to data collection and taxonomic identification. Writing of the manuscript was led by ARG with input from all co-authors.

## DATA AVAILABILITY

Genera distribution data summary and GenBank accession numbers are provided in supplementary table 1. Plot location information openly available at www.andesconservation.org

## Acknowledgements

This paper is a product of the Andes Biodiversity and Ecosystem Research Group consortium. We thank SERFOR, SERNANP and personnel of Manu National Park for logistical assistance and permission to work in the protected area and the buffer zone. We thank Ricardo Segovia for compilation of many GenBank sequences and helpful advice. We thank the Gordon and Betty Moore Foundation Andes to Amazon program and the Blue Moon Fund for support for the plot network. ARG. is supported by a NERC-E3DTP studentship (grant number: NERC NE/L002558/1). PM was supported by NERC (NE/G018278/1) and ARC (DP170104091). KJF is supported by the US National Science Foundation (NSF DEB LTREB 1754664). MS was supported by NSF DEB LTREB 1754647. We also thank two anonymous reviewers for their suggestions and reflections.

